# The Ccr4-Not complex monitors the translating ribosome for codon optimality

**DOI:** 10.1101/854810

**Authors:** Robert Buschauer, Yoshitaka Matsuo, Ying-Hsin Chen, Najwa Alhusaini, Thomas Sweet, Takato Sugiyama, Ken Ikeuchi, Jingdong Cheng, Yasuko Matsuki, Andrea Gilmozzi, Otto Berninghausen, Thomas Becker, Jeff Coller, Toshifumi Inada, Roland Beckmann

**Affiliations:** Gene Center and Center for Integrated Protein Science Munich, Department of Biochemistry, Feodor-Lynen-Str. 25, University of Munich, 81377 Munich, Germany; Graduate School of Pharmaceutical Sciences, Tohoku University, Sendai 980-8578, Japan; Center for RNA Science and Therapeutics, Case Western Reserve University, School of Medicine, 10900 Euclid Ave., Cleveland OH 44106-4960, USA

## Abstract

Control of mRNA decay rate is intimately connected to translation elongation but the spatial coordination of these events is poorly understood. The Ccr4-Not complex initiates mRNA decay through deadenylation and activation of decapping. Using a combination of cryo-electron microscopy, ribosome profiling and mRNA stability assays we show recruitment of Ccr4-Not to the ribosome via specific interaction of the Not5 subunit with the ribosomal E-site. This interaction only occurs when the ribosome lacks accommodated A-site tRNA, indicative of low codon optimality. Loss of Not5 results in the inability of the mRNA degradation machinery to sense codon optimality. Our analysis elucidates a physical link between the Ccr4-Not complex and the ribosome providing mechanistic insight into the coupling of decoding efficiency with mRNA stability.

---

The Ccr4-Not (carbon catabolite repressor 4 -negative on TATA) complex is an essential and conserved multisubunit complex which comprises at least six core subunits arranged in a modular architecture. While Ccr4-Not has roles in almost every aspect of regulating gene expression (i.e. chromatin remodeling, transcription, mRNA export, and miRNA regulation) its most defined and best understood role is in mRNA degradation, serving as the major cytoplasmic deadenylase besides Pan2-Pan3 (*1, 2*). Two subunits of Ccr4-Not (Caf1 and Ccr4) are poly(A)-specific nucleases that initiate decay of the majority of mRNAs. Their activity in shortening the 3’-poly-A-tail is typically followed by removal of the 7-methylguanylate cap at the 5’-end by Dcp2-Dcp1 holoenzyme and then subsequent 5’-to-3’ degradation by Xrn1 (*3, 4*). For most transcripts, decapping is strictly dependent on prior deadenylation. However, little is known about the spatial organization allowing the coordination of deadenylation with decapping by Ccr4-Not.

The control of mRNA stability is a critical process with the rates of mRNA decay setting the overall level of gene expression and the final concentration of protein product. Half-lives of mRNAs vary greatly between transcripts, and codon optimality has been established as a major parameter determining mRNA half-life (*5*). The timely decay of short-lived mRNAs containing non-optimal codons involves the Ccr4-Not complex (*6*) and the Ccr4-Not associated helicase and activator of decapping Dhh1 (*7*). Recent RNA-binding studies showed enrichment of Dhh1 and other decapping enhancers on non-optimal mRNAs on a genome wide scale (*8*). The molecular principles underlying monitoring of codon optimality during translation, however, have remained enigmatic. Since non-optimal codons affect the decoding kinetics and mRNA degradation occurs co-translationally, it is expected that codon optimality is directly monitored on the ribosome. A physical link of the Ccr4-Not complex to the ribosome has been suggested previously, in particular to polyribosomes (*9, 10*). The Not4 subunit of the complex, an E3-ligase, ubiquitinates proteins of the 40S small ribosomal subunit (*11, 12*), and the Not5 subunit appears to be important for mRNA translatability and polyribosome levels in yeast (*13*). We therefore aimed at shedding light on the interplay between Ccr4-Not and the translation machinery in the context of mRNA homeostasis.

## The Not5 subunit anchors Ccr4-Not to the ribosome

To characterize a physical link between the ribosome and Ccr4-Not, we co-purified endogenous ribosome bound Ccr4-Not complexes from *Saccharomyces cerevisiae* using tagged Not4 (a stable subunit of Ccr4-Not) as bait. We performed pull-downs following sucrose density gradient centrifugation and subsequent isolation of a monosome 80S and a polysome fraction, both stabilized by the antibiotic tigecycline to prevent ribosomal run-off. Single particle cryo-EM analysis of the monosome fraction resulted in a structure of an 80S ribosome with a vacant A-site, a tRNA in the P-site and additional mostly alpha helical density in the E-site (Fig. 1A). The good overall resolution of 2.8 Å (fig S1, A-D) allowed us to identify the extra density as the N-terminal domain (NTD) of Not5 (residues 2-113). Not5 is a highly conserved component of Ccr4-Not (CNOT3 in human) with known roles in controlling mRNA stability (*14, 15*). We could build an atomic model of the previously structurally uncharacterized Not5-NTD (Fig. 1B). The remaining part of Not5 and also the entire Ccr4-Not complex were apparently flexibly linked to the NTD and therefore not resolved, with exception of a small helical bundle extending from the NTD near the 40S head, which we could see but not model in a chemically cross-linked cryo-EM sample (fig. S1, E and F). Yet, presence of the entire complex in the cryo-EM sample was confirmed by SDS-PAGE and mass-spectrometry (Fig. 1C). Thus, anchoring of the complex to the E-site by the Not5 subunit still allows for a high degree of spatial freedom for the complex with respect to the ribosome.

**Fig. 1.**
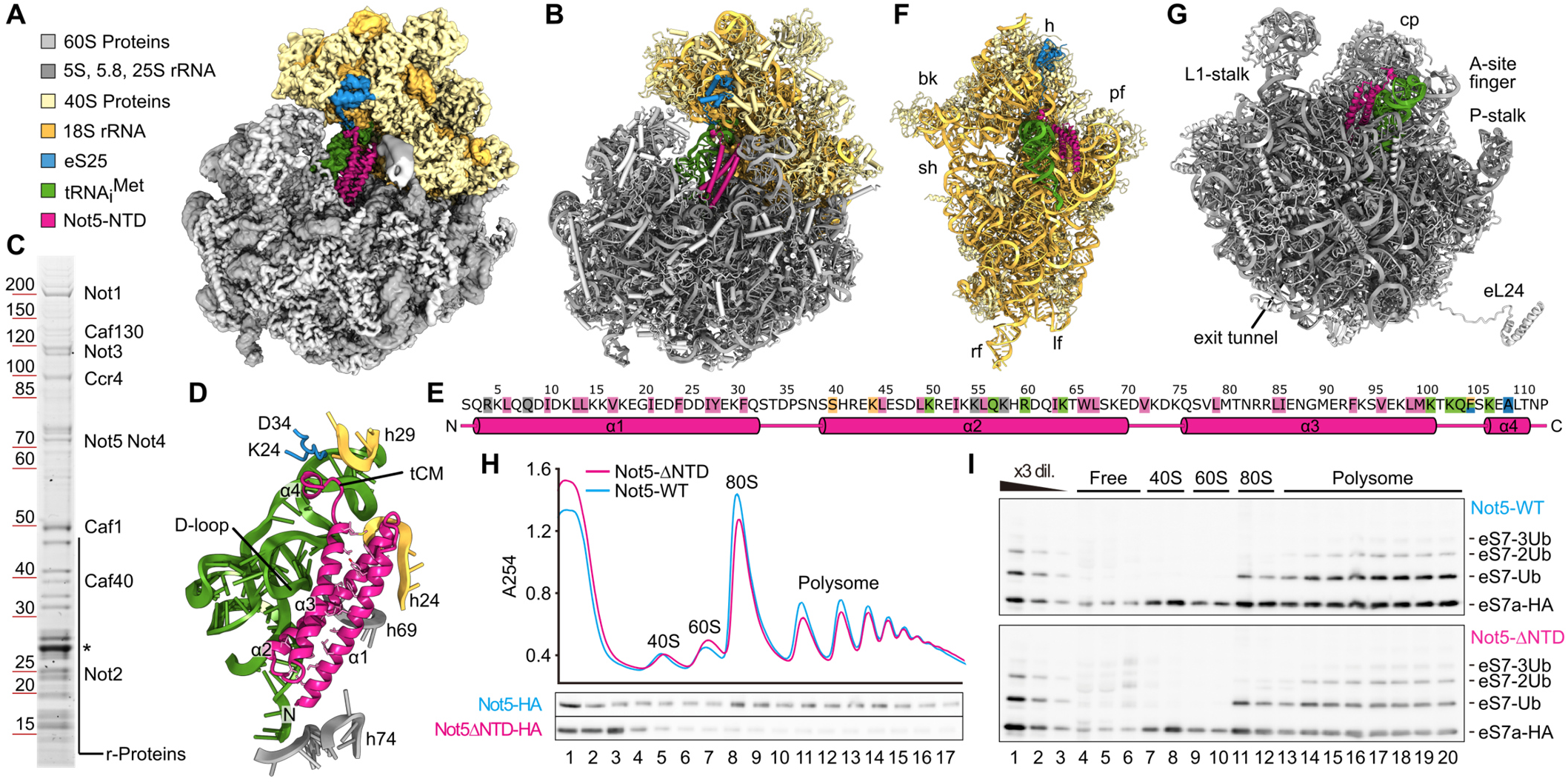
The N-terminus of Not5 binds to the ribosomal E-site. (**A**) Cryo-EM density map of the Not5-80S complex. (**B**) Atomic model of the Not5-80S complex. For visibility of Not5-NTD, protein uL1 is not displayed (**C**) SDS-PAGE analysis of the Not4 affinity purification. The core subunits of the Ccr4-Not complex as well as 80S ribosomes were co-purified, as confirmed by mass-spectrometric analysis. The asterisk indicates the band corresponding to TEV-protease. (**D**) Overview of Not5-NTD structure and orientation with respect to interacting ribosomal features and tRNA_i_^Met^. The tRNA-clamp-motif (tCM) of Not5-NTD is indicated. Residues that establish the hydrophobic core of the three-helix bundle are shown as sticks. (**E**) Topology of Not5-NTD. Important residues are highlighted (pink: hydrophobic core, green: interaction with tRNA, blue: interaction with eS25, beige: interaction with 18S rRNA, grey: interaction with 25S rRNA). (**F**) Atomic model of the 40S subunit, P-site tRNA and Not5-NTD, as seen from the ribosomal inter subunit space. Locations of head (h), beak (bk), platform (pf), shoulder (sh), left foot (lf) and right foot (rf) of the 40S subunit are indicated. (**G**) Atomic model of the 60S subunit, P-site tRNA and Not5^2-113^, as seen from the ribosomal inter subunit space. Location of the central protuberance (cp) as well as other characteristic features of the 60S subunit are indicated. (**H**) Density gradient profiles of Not5-HA and Not5-∆NTD-HA strains and corresponding western blot detection of the respective Not5 proteins. (**I**) Western blot detection of ribosomal protein eS7-HA in the fractions of a density gradient from an eS7-HA strain.

The NTD of Not5 comprises a three-α-helix-bundle (α1: 3-32, α2: 39-70, α3: 76-101) and a short linker followed by a β-turn (103-106), which directly leads into a short fourth α-helix (α4: 107-111) (Fig. 1, D and E). The three-helix bundle establishes a stable hydrophobic core and the entire NTD binds to the ribosome in the E-site by precisely spanning the distance between the 60S and the 40S ribosomal subunit. The domain directly interacts with the P-site tRNA, the 25S rRNA, 18S rRNA and the N-terminal tail of the ribosomal protein eS25, suggesting that binding requires a fully assembled 80S ribosome with tRNA in the P-site (Fig. 1, E-G). In order to rule out the possibility of an artifact induced by tigecycline, we repeated structure determination in absence of antibiotics and confirmed the presence of the NTD in the same position and conformation (fig. S1, G-I). Moreover, the same overall architecture was also observed for the complexes after chemical cross-linking or when isolated from the polysome fraction (see below).

To test whether the NTD is required for ribosome association of the Ccr4-Not complex *in vivo* we generated a yeast strain carrying a Not5 construct lacking the NTD: analysis of this mutant using sucrose density gradient centrifugation revealed that association of the Ccr4-Not complex with (poly)-ribosomes was decreased as evident from the signal for the Not5 subunit (Fig. 1H and fig. S2). Most of the mutant complex was observed in the top fraction of the gradients with some signal remaining throughout the gradient. We therefore conclude that the NTD of Not5 makes a critical contribution, which is required to facilitate the association of the Ccr4-Not complex with the ribosome, in particular the polysome.

The remaining association with the translation machinery is likely to reflect a second binding mode of the Ccr4-Not complex that could involve the interaction of the Not4 subunit with ribosomal proteins for ubiquitylation. Consistent with this idea, the modification of eS7 still occurred in the Not5-∆NTD strain (Fig. 1I and fig. S2).

## Not5 targets initiating and elongating ribosomes

We initially used the native Ccr4-Not:80S complexes from the monosome fraction of the sucrose gradient and expected to find a mixture of different mRNAs and tRNAs. To our surprise, however, the codon present within the P-site was well resolved and identified as AUG (Fig. 2A). Moreover, the tRNA could be unambiguously identified as the corresponding initiator tRNA_i_^Met^, and not the elongator tRNA^Met^ (Fig. 2B). These data suggest an association of the Ccr4-Not complex with late initiation complexes. To confirm this, we probed by Northern blotting for individual tRNAs in both, total 80S fractions and Not4 pull-down monosome fractions and in presence or absence of tigecycline or cycloheximide (Fig. 2C). As expected, we observed a strong enrichment of tRNA_i_^Met^ and not tRNA^Ala^ (as control) in the ribosomal samples upon pull-down independent of tigecycline, whereas co-purifications in presence of cycloheximide yielded significantly less ribosomes. A structural feature of tRNA_i_^Met^ compared to some other tRNAs is a short D-loop (*16*), which in our structure touches the three-helix-bundle of Not5 (Fig. 2, B and D). This may suggest some degree of specificity of Not5 conferred by proximity and probing of the tRNA D-loop. Finally, interaction with late initiation complexes was also consistent with the absence of density for a nascent polypeptide chain (Fig. 2I) and with the results of selective ribosome profiling following co-immunoprecipitation using Not4 as a bait or the ribosomal protein uL30 as a control (Fig. 2E): In comparison to the control, we observed a two-fold enrichment of ribosomes on the initiation codon across all messages (Fig. 2F). A functional role of the Ccr4-Not complex in translation initiation was suggested previously, based on the observation that polysome levels decrease upon deletion of Not5 (*12*). Consistently, also our polysome profiles displayed a small but significant difference between wildtype and Not5-∆NTD cells with respect to amounts of active ribosomes (80S and polysomes) versus free ribosomal subunits (Fig. 1H and fig. S2).

**Fig. 2.**
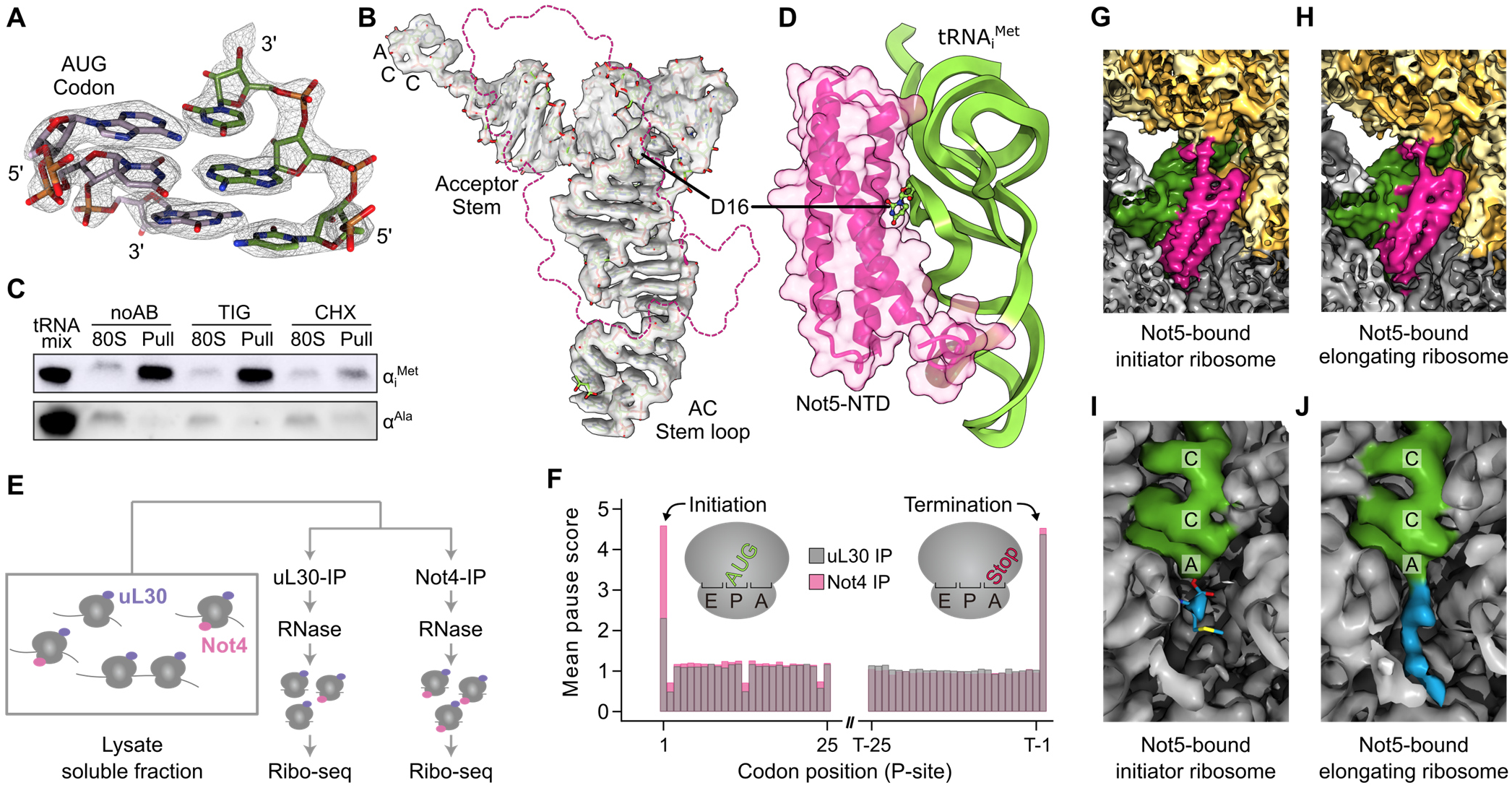
Not5 interacts with initiating and elongating ribosomes. (**A**) Cryo-EM density and atomic model of the codon-anticodon interaction. (**B**) Cryo-EM density and atomic model of tRNA_i_^Met^. The position of Not5-NTD is outlined in pink. (**C**) Northern blot against tRNA_i_^Met^ (top) and tRNA^Ala^ (bottom). Same amounts of ribosomes were loaded before (80S) and after affinity purification (Pull). The samples were prepared without antibiotics (noAB), in presence of tigecycline (TIG), or in presence of cycloheximide (CHX). (**D**) D-loop recognition through Not5. Not5-NTD (pink) is contacting the D-loop of tRNA_i_^Met^ (green). (**E** to **F**) Selective ribosome profiling of Ccr4-Not associated ribosomes (Not4 IP) and total ribosomes (uL30 IP). Not4 bound ribosomes are enriched on the initiation codon. (**G** to **H**) Cryo-EM densities of the Not5 binding site in the initiator and elongating ribosome. Structures were lowpass filtered to 5 Å. (**I**) Peptidyl transferase center (PTC) of the Not5-80S complex. There is no density corresponding to a nascent poly peptide chain extending beyond the initial methionine (blue). (**J**) PTC of the elongating Not5 bound ribosome. There is clear density for a nascent poly peptide chain (blue).

We next asked whether the Ccr4-Not complex also binds to elongating ribosomes and determined cryo-EM structures from heavy fractions of a sucrose gradient (between 3 and 6 ribosomes). Also here, we observed a population of ribosomes with Not5 bound in the E-site, tRNA present in the P-site and a vacant A-site. Altogether this structure resembled our previous observations with Not5 bound to the late initiation complex ribosome (Fig. 2, G and H). However, the local resolution of the tRNA was not high enough as to unambiguously identify the tRNA, suggesting the presence of a mix of different tRNAs species. Yet, the shape of the D-loop was very similar to that of tRNA_i_^Met^ present in the late initiation complex, supporting specificity of Not5 for D-loop shape. In contrast to the initiation structure, there was clear density for a nascent chain extending from the peptidyl transferase center into the ribosomal exit tunnel, confirming the elongating state of the ribosomes (Fig. 2, I and J).

Taken together, these data show that Not5 links the Ccr4-Not complex to both, late initiating and also elongating ribosomes within the open reading frame.

## Molecular interactions of the Not5 NTD

A strong interaction of the NTD with the ribosome is established through the P-site tRNA. It involves multiple hydrogen bonds between the backbone of A14-D16 of the tRNA D-loop with residues Q57, R60 and K64 of helix α2 of Not5 (Fig. 3A), whereas the backbone of residues G22 and C23 of the D-arm hydrogen bonds with K50 of α2 and K101 of α3 (Fig. 3, B and C). Another tight interaction involves residues K103 – L110 of Not5, which we termed the tRNA clamp-motif (tCM) (Fig. 3D) The phosphate backbone of residues U42 and G43 of the tRNA anticodon stem loop is hydrogen bonded by the side chain of K103 and the peptide backbone nitrogens of residues Q104, F105 and K107. Notably, this interaction is further stabilized by the N-terminal tail of the ribosomal protein eS25 (Fig. 3, D and E). This tail of eS25 is flexible and usually not well resolved in cryo-EM or crystal structures of eukaryotic ribosomes. In the presence of Not5, however, it is stabilized and can be observed extending from the globular part of eS25 at the head of the small ribosomal subunit to the ribosomal P-site (Fig. 1E and 3C). There, residues K25 and K29 of eS25 form hydrogen bonds with the carbonyl atoms of F105 and A109 of Not5, respectively. Thereby, eS25 holds the tCM of Not5 in place and, together eS25, tCM and 18S rRNA form a groove which tightly associates with the tRNA backbone (Fig. 3E). Residue W27 of eS25 pins the flexible N-terminal tail to this location through a stacking interaction with G1575 of the 18S rRNA between h29 and h42. The same rRNA base also stabilizes the β-turn of Not5 through a stacking interaction between the ribose and the sidechain of F105 of Not5 (Fig. 3D). Notably, K25, W27 and K29 of eS25 belong to the highly conserved KKKWSK motif and a human K33E mutation (corresponding to yeast K25E) has been found in thyroid carcinoma cells (*17*). The intricate interactions between tRNA, eS25 and 18S rRNA tightly fix the conformation of the tCM. The central residue F105 of the tCM is coordinated by rRNA through the sidechain, by eS25 through the carbonyl oxygen and by tRNA through the backbone nitrogen. Additional interactions of the Not5 NTD with the 18S rRNA backbone phosphates of G999 and G997 of h24 is established by S40 and K44 of Not5, whereas H41 of Not5 interacts with the ribose of A907 of rRNA h23 (Fig. 3F). Interaction of Not5 with the 25S rRNA is established by hydrogen bonding of K55 and K58 of α2 with the backbone phosphates of U2269 and U2268 in the vicinity of rRNA helix h69 (Fig. 3G). A second interaction with 25S rRNA involves helix h74 and helix α1 of Not5 (Fig. 3H). Here, residue R4 of Not5 stacks on the ribose of A2419 while forming a hydrogen bond with the phosphate of A2804, and Q8 stacks and hydrogen bonds with the base of G2418.

**Fig. 3.**
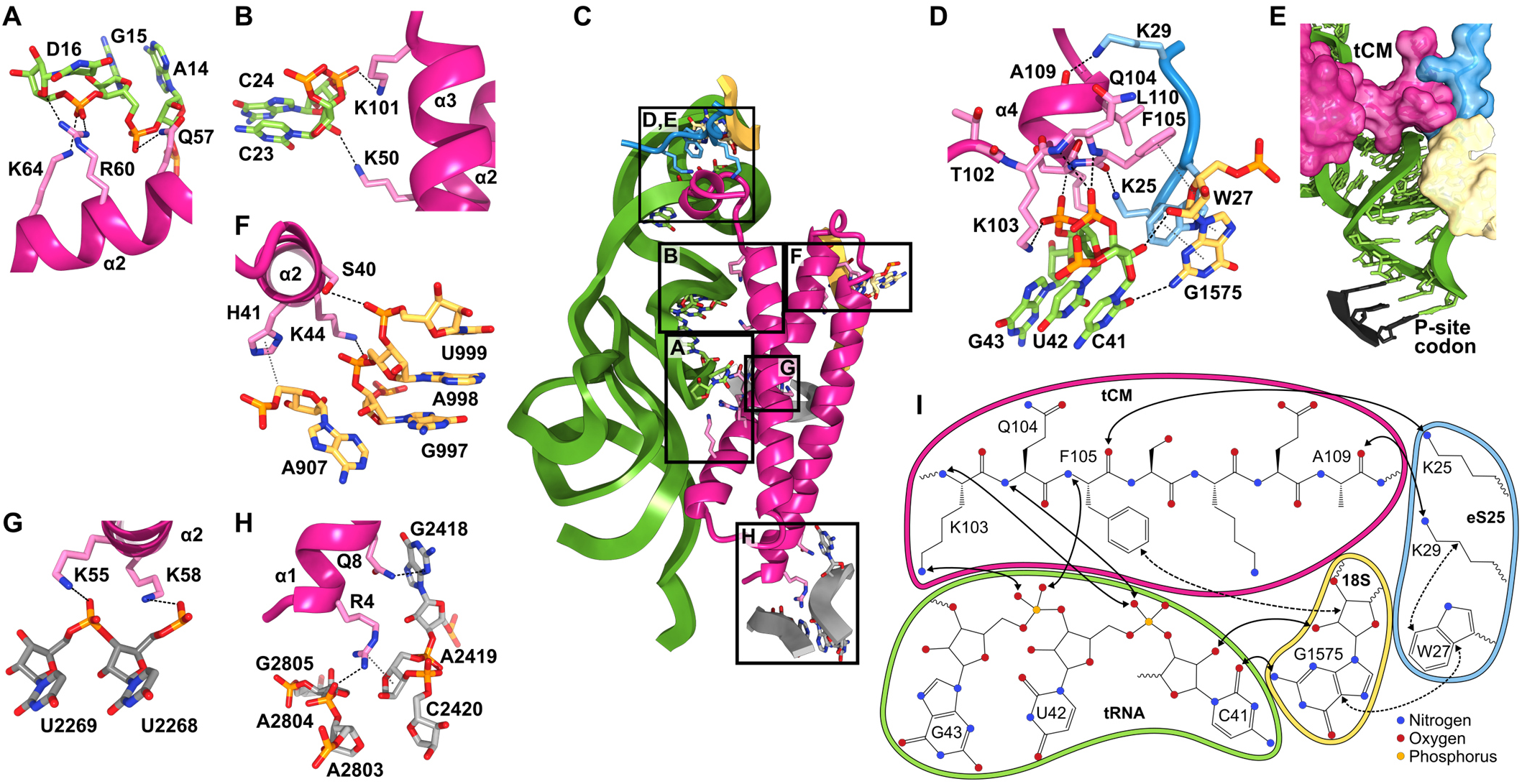
Not5-80S complex interface. (**A**) Interactions of Not5-NTD with the D-loop of tRNA_i_^Met^. Hydrogen-bonds and ion bridges are shown as black dashed lines. (**B**) Interactions of Not5 with the D-stem of tRNA_i_^Met^. (**C**) Overview of the interaction sites of Not5-NTD and the initiator-80S ribosome. The interaction sites shown in more detail in the other panels are indicated. Interacting residues are shown in stick representation. (**D**) Intricate interactions between the tRNA-clamp-motif (tCM) of Not5, the anticodon stem loop of tRNA_i_^Met^, the N-terminus of eS25 and 18S rRNA. (**E**) Surface representation of tCM, eS25 and 18S rRNA and atomic model of the anticodon stem loop of tRNA_i_^Met^. Hydrogen-bonds and ion bridges are shown as black dashed lines. Hydrophobic interactions are shown in grey dotted lines. (**F**) Interactions of Not5-NTD with 18S rRNA. (**G** to **H**) Interactions of Not5 with 25S rRNA. (**I**) Schematic summary of the binding interface of the tCM of Not5 with the initiatior 80S. Hydrogen bonds and ion bridges are indicated with solid arrows; hydrophobic interactions are shown as dashed arrows.

Taken together, the Not5 NTD engages in a highly specific and complex binding mode in the ribosomal E-site involving 40S and 60S ribosomal rRNA as well as eS25 and a P-site tRNA with a small D-loop.

Notably, the human homolog of Not5, CNOT3, contains a highly conserved NTD suggesting a conserved function of CNOT3 as an anchor to the human ribosome. Interestingly, multiple cancer mutations cluster in the NTD of CNOT3 (*17*) with one of the most frequent mutations being R57W/E (K58 in yeast) (*18*). K58 directly interacts with the phosphate backbone of 25S rRNA in our structure (Fig. 3G) and an analogous interaction can be predicted for R57 of the human homolog. Mutation of K58 (or R57 in human) to tryptophan or glutamate would not allow hydrogen bond formation and therefore destabilize the interaction.

## Not5 competes with eIF5A dependent on A-site occupation

We observed Not5 binding to ribosomes in the post-translocation state, in which ribosomes usually carry a deacylated tRNA in the E-site, a peptidyl-tRNA in the P-site (--/PP/EE), but no A-site tRNA. Not5 can apparently bind to this post state conformation as soon as both E-site and A-site are vacant (--/PP/--) (Fig. 4A). In this state, the small ribosomal subunit is rotated towards the E-site and the L1-stalk cannot adopt the far inside conformation due to a potential clash with the 40S subunit (fig. S3). In contrast, upon accommodation of the A-site tRNA (AA/PP/--), the 40S ribosomal subunit undergoes a conformational change in the opposite direction - known as subunit rolling - in combination with repositioning of the head (*19*) (Fig. 4B). Subunit rolling is visualized by superposition of the 40S subunits of the two states upon alignment of the 60S subunits (Fig. 4C). Together, these movements result in characteristic remodeling of the E-site and allow, for example, the far inside conformation of the L1 stalk, which is necessary to accommodate eIF5A. In the presence of cycloheximide we observed enrichment of this pre-translocation state with A- and P-site tRNAs in different preparations, in which we also found eIF5A bound to the E-site (*20*).

**Fig. 4.**
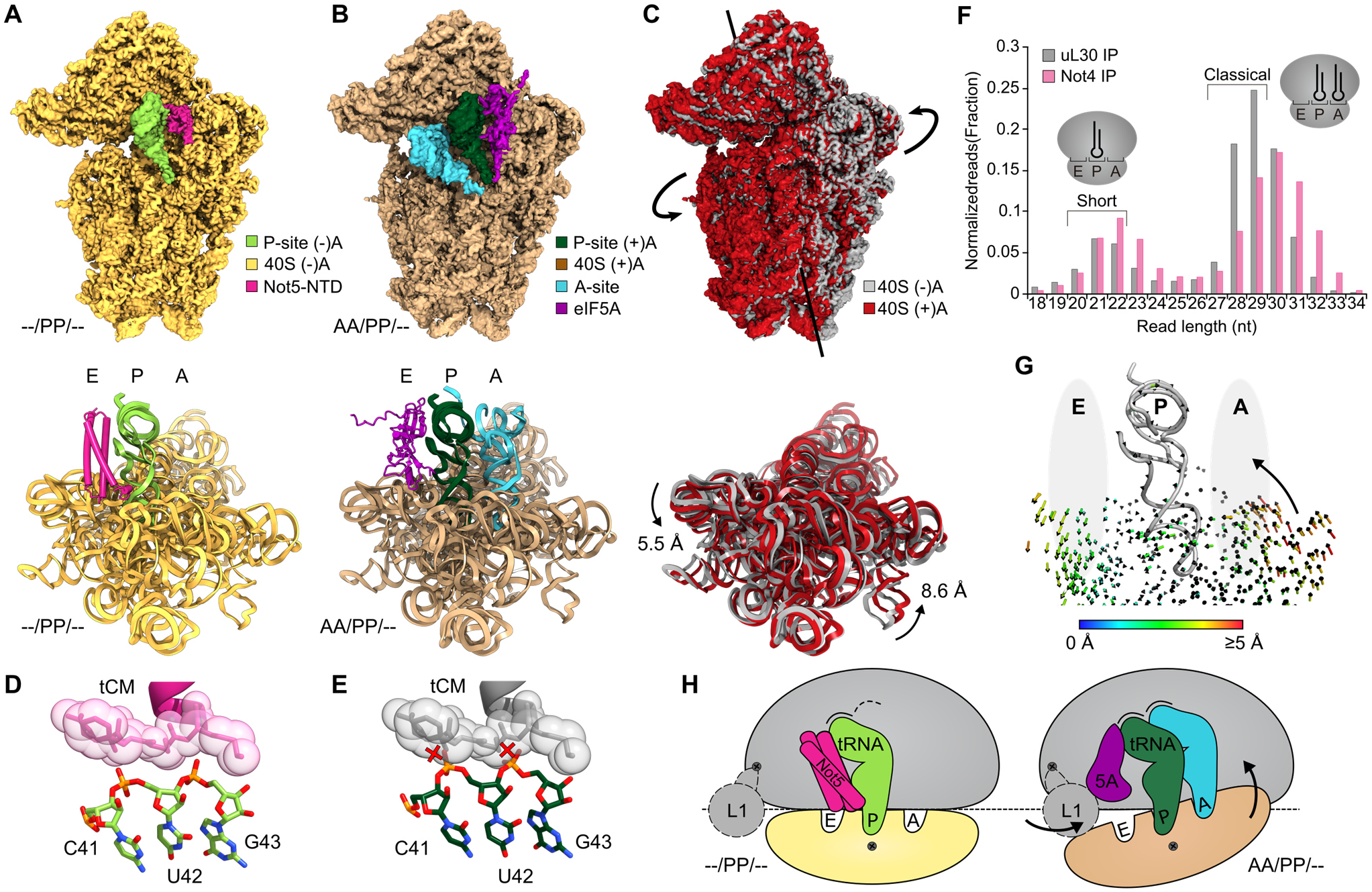
eIF5A and Not5 compete for the E-site in an A-site dependent manner. (**A**) Density (top) and model (bottom) of the Not5-bound 40S subunit of the (--/PP/--) 80S ribosome. Model is viewed from the head. For simplicity, only 18S rRNA is shown. (**B**) Density (top) and model (bottom) of the eIF5A-bound 40S subunit of the (AA/PP/--) 80S ribosome. (**C**) Superposition of the densities (top) and the models (bottom) of the 40S subunits of the (--/PP/--) and the (AA/PP/--) ribosomes. Subunit rolling is indicated. The structures in a, b and c were aligned with respect to the 60S subunits. (**D**) The tCM of Not5 is closely contacted by the anticodon stem of the P-site tRNA in the (--/PP/--) situation. (**E**) The conformational changes of the P-site tRNA upon A-site accommodation would cause clashes between tRNA backbone and the tCM (red crosses). (**F**) Length distribution of mRNA fragments during selective ribosome profiling using Not4 and uL30 (total ribosomes) as baits. (**G**) Illustration of A-site E-site coupling. The small arrows represent quantified trajectories of Cα atoms (Protein) and P atoms (RNA) necessary for transition between (--/PP/--) and (AA/PP/--) conformations. (**H**) Schematic summary of subunit rolling and the competition between eIF5A and Not5.

Not5 and eIF5A binding to a vacant E-site appears to be mutually exclusive, not only because of a direct steric clash between the two factors, but rather because of different conformational requirements for E-site binding. Comparison of the Not5-bound (--/PP/--) ribosome with the eIF5A-bound (AA/PP/--) ribosome shows that upon A-site accommodation, the anticodon stem loop of the P-site tRNA has moved towards the E-site. This involves nucleotides 41-43, which are clamped by the tCM of Not5 and this subtle movement of the tRNA backbone (~1.6 Å) would cause clashes with the tCM of Not5 upon A-site accommodation (Fig. 4, D and E). This explains why the NTD of Not5 can act like a precise tape measure which can stably and specifically interact with the ribosomal E-site in the absence of an accommodated A-site tRNA. In agreement with this, we only observe small populations of Ccr4-Not:80S complexes in the presence of cycloheximide, which stabilizes the unfavored pre-translocation state with an A-site tRNA.

Moreover, preferential association in the presence of empty E- and A-sites was further confirmed by selective ribosome profiling using Not4 as bait: we observed enrichment of short mRNA fragments (~21mers), which have previously been shown to be derived from ribosomes with an empty A-site (*21*) (Fig. 4F). The allosteric coupling between A- and E-site of the 80S ribosome is illustrated by quantified trajectories of ribosomal proteins (Cα-atoms) and RNA (P-atoms) that describe a transition between the Not5-bound post state and the A-site tRNA accommodated pre state (Fig. 4G).

In summary, we found that the NTD of Not5 can probe the ribosomal E-site in a highly selective way resulting in the Ccr4-Not complex most efficiently interacting with ribosomes with a vacant A-site, as schematically summarized in Fig. 4H. As a result, a coherent picture emerges in which the Ccr4-Not complex and eIF5a facilitate a differential read out of the translational state of the ribosome via E-site probing: the Ccr4-Not complex would be recruited with preference to ribosomes displaying low decoding efficiency and thereby increase the likelihood of mRNA deadenylation and degradation (*22*). In contrast, the competing eIF5A would preferentially interact with ribosomes remaining longer in the pre-translocation state because of slow peptidyl transferase kinetics, thereby allowing for stabilization of a productive peptidyl transferase center geometry and effectively counteracting Ccr4-Not recruitment.

## Not5 and optimality guided mRNA degradation

Codon optimality is a major determinant of mRNA degradation with the proportion of non-optimal codons within the transcript determining its half-life (*5*). In essence, a non-optimal codon is one whose functional tRNA concentration is low, resulting in prolonged vacancy of the ribosomal A-site. Since eviction of the E-site tRNA does not require A-site accommodation (*23*), prolonged vacancy of the A-site results in an increased probability of a ribosome adopting the (--/PP/--) state, which can be recognized by Not5. Thus, a ribosome with an empty E-site in the post-state conformation could be specifically sensed by Not5 as a proxy for the presence of a non-optimal codon in the A-site. To test this hypothesis, we performed selective ribosome profiling upon co-immunoprecipitation of Ccr4-Not:ribosome complexes, again using Not4 as bait. The data revealed a significant correlation between non-optimality of the A-site codon and presence of Ccr4-Not on the translation machinery (Fig. 5A). The A-site codons with the strongest correlation were CGG, CGA, CTG and AGT, which are among the most non-optimal codons (*5*) (Fig. 5B).

**Figure 5:**
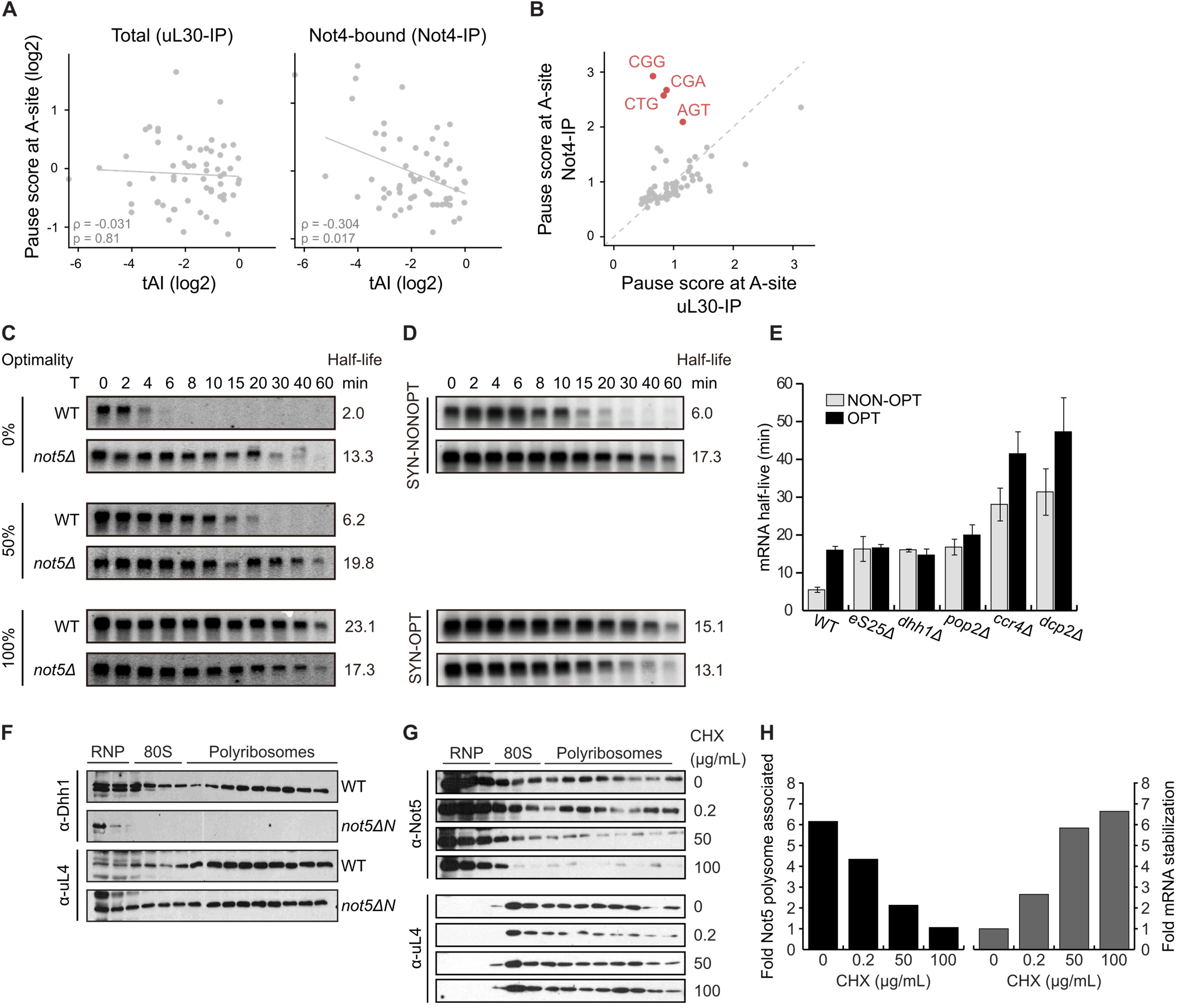
The Not module and eS25 are involved in decapping of non-optimal mRNAs. (**A**) Selective ribosome profiling of Ccr4-Not associated ribosomes (Not4-IP) and total ribosomes (uL30-IP). Presence of Not4 on the ribosome is correlated with non-optimal codons in the A-site (based on tRNA adaptation index, tAI). (**B**) Enrichment of specific non-optimal codons in the A-site of Ccr4-Not associated ribosomes as determined by selective ribosome profiling. (**C**) Half-life determination of HIS3 reporter mRNA through northern blotting at various timepoints after transcriptional shutoff (T) in either wildtype or *not5∆* strains. (**D**) Half-life determination of optimal and non-optimal PGK1pG reporter mRNAs through northern blotting at various timepoints after transcriptional shutoff (T) in either wildtype or *not5∆* strains. (**E**) Half-lives of optimal and non-optimal PGK1pG reporter mRNAs in wildtype and *eS25∆*, *dhh1∆*, *pop2∆*, *ccr4∆* and *dcp2∆*, as determined by northern blotting. (**F**) Presence of Dhh1 (or uL4 as control) on the translation machinery determined by western blotting of sucrose density gradients in wildtype and *not5∆NTD* strains. (**G**) Presence of Not5 (or uL4 as control) on the translation machinery determined by western blotting of sucrose density gradients in presence of different concentrations of cycloheximide. (**H**) Quantification of the association of Not5 with the translation machinery and fold stabilization of a non-optimal PGK1pG reporter mRNA in dependence of cycloheximide concentration.

Next, we wanted to test whether specifically the Not5 subunit of the Ccr4-Not complex is involved in optimality dependent mRNA degradation and evaluated the decay rates of reporters previously developed to monitor codon optimality. We observed that loss of Not5 indeed stabilized non-optimal mRNAs, apparently disabling cells to degrade mRNAs according to codon usage. Specifically, the half-life of non-optimal mRNAs in absence of Not5 was equal to that of optimal mRNAs in WT cells (Fig. 5, C and D). These data demonstrate that Not5 is required by the cell to communicate codon optimality to the mRNA degradation machinery, and are in agreement with our hypothesis of codon optimality sensing by E-site probing through Not5. Curiously, deletion of the Not5 NTD alone yielded inconsistent results (data not shown) and hint at a second layer of regulatory function for this domain. Consistent results, however, have been reported for the mammalian system, in which deletion of the NTD of CNOT3 led to an overall stabilizing effect for several mRNAs (*24*).

Moreover, in agreement with the interaction of the NTD of Not5 with the ribosomal protein eS25 as observed in our structure, deletion of both copies of eS25 also resulted in the mRNA degradation machinery’s inability to sense codon optimality (Fig. 5E).

Since it has previously been demonstrated that the small helicase Dhh1 is required for decapping of non-optimal mRNAs and because Dhh1 associates with Ccr4-Not (*7*), we determined whether the Not5/E-site interaction is required for presence of Dhh1 on the translation machinery. We found that deletion of the NTD of Not5 was indeed sufficient to remove Dhh1 from (poly)-ribosomes (Fig. 5F).

Lastly, we evaluated if Not5 can be chased from ribosomes through cycloheximide and whether this also causes a stabilization of non-optimal mRNAs. As described above, cycloheximide stabilizes the ribosome in the (AA/PP/--) state, which is conformationally unfavorable for Not5 binding to the E-site. As expected, treatment of cells with cycloheximide resulted in a dose-dependent depletion of the Ccr4-Not complex from polyribosomes (Fig. 5G) and, importantly, the level of Not5 depletion from (poly)-ribosomes was matched by an equal fold stabilization of non-optimal mRNAs (Fig. 5H). Similar mRNA stabilizing effects of cycloheximide were observed before, however, rather attributed to an overall protective property of mRNA-bound ribosomes (*25*). Here we suggest that E-site antibiotics such as cycloheximide disrupt the association of Not5 with the ribosome and hence result in an alteration of mRNA decay kinetics.

## Discussion

Taken together, we discovered a physical link between the Ccr4-Not complex and the ribosome mediated by the Not5 subunit and, in addition, we found that Ccr4-Not association to the ribosome correlates with non-optimal codons in the A-site. Strikingly, the specificity for ribosomes in the post-state with simultaneous vacancy of ribosomal E- and A-sites provides the molecular basis for a sensing mechanism for codon optimality by the Ccr4-Not complex. It also explains how the presence of non-optimal codons can be communicated to the mRNA degradation machinery, thereby shedding light on the coupling between mRNA translation and decay. The observed recruitment of the Ccr4-Not complex to elongating ribosomes with non-optimal codons can therefore be rationalized by the subsequent triggering of mRNA decay. This is in contrast to the consequence of the competing interaction of eIF5a with the E-site that binds in the presence of an A-site tRNA and supports PTC activity in order to continue elongation.

It is curious, however, that we also observe a large fraction of the Ccr4-Not complex bound to late initiation 80S complexes. From a structural perspective these complexes are ideal targets for Ccr4-Not, because they are fully assembled 80S ribosomes in the post state with tRNA in the P-site and empty A- and E-sites. With respect to the deadenylation function of Ccr4-Not, however, especially in context of codon-optimality, the location seems puzzling. However, one possibility is that independent of codon optimality, mRNA decay could be triggered in this case due to stalling of translation initiation. This is supported by our finding that these initiation complexes were mainly present in the 80S monosome and not in the polysome fractions. Another possibility is that we were observing a later stage in the mRNA degradation pathway: Considering the closed loop mRNA model (*26, 27*), one could speculate that Ccr4-Not is handed over from the 3’ end to the 5’ end of the mRNA subsequent to deadenylation. Such a handover could be beneficial since anchoring the complex to the E-site of an 80S ribosome located on the start codon would be an ideal way to assemble and activate the decapping machinery within spatial proximity to the 5’ cap, while probably inhibiting further initiation. In line with this idea, recent RNA-binding studies of deadenylation and decapping factors showed an interesting distributed allocation on the 5’- and 3’-ends of the transcripts (*8*). Since the functionality of Ccr4-Not extends far beyond mRNA degradation, binding to late initiation complexes could also be involved in entirely different processes such as co-translational assembly of multi-subunit complexes as recently suggested for SAGA complex and proteasome (*28, 29*). The diverse activities of the truly multifunctional Ccr4-Not complex and its relationship to the translation machinery will be further dissected in future studies. Understanding the molecular details of this relationship will be crucial for comprehension of Ccr4-Not function in health and disease.

## Materials and Methods

### Yeast strains and genetic methods

Gene disruption and C-terminal tagging were performed by established recombination techniques as previously described (*30, 31*). The *S. cerevisiae* strains used in this study are listed in Table S1.

### Plasmids construct

All recombinant DNA techniques were performed according to standard procedures using *E. coli* DH5a for cloning and plasmid propagation. All cloned DNA fragments generated by PCR amplification were verified by sequencing. Plasmids used in this study are listed in Table S2.

### Native complex purification

A *S. cerevisiae* strain with genomically tagged Not4 (Not4-FTPA) was cultured in 10 l YPD medium supplemented with 5 µg/ml ampicillin and 10 µg/ml tetracycline and harvested by centrifugation at OD_600_ of 0.9. The pellet was washed in water and lysis buffer (20 mM HEPES pH 7.4, 100 mM KOAc, 10 mM Mg(OAc)_2_, 1 mM DTT, 0.5 mM PMSF, 100 µg/ml tigecycline, protease inhibitor cocktail tablet (Roche)) and frozen in liquid nitrogen. Lysis was performed by grinding the frozen cell pellet in a Freezer Mill (6970 EFM). The ground powder was thawed in 15 ml lysis buffer and the lysate was cleared by centrifugation at 12000 x g for 15 min at 4 °C. Cleared lysate was distributed and layered on top of 6 sucrose density gradients (10-50% sucrose in 20 mM HEPES pH 7.4, 100 mM KOAc, 10 mM Mg(OAc)_2_, 1 mM DTT, 10 µg/ml tigecycline in 25 × 89 mm polyallomer tubes, SW32, Beckman Coulter) and centrifuged at 125755 x g for 3 h at 4 °C. The gradients were subsequently fractionated from top to bottom using a Gradient Master (BioComp). The fractions corresponding to the 80S peak (monosome sample) or corresponding to polysomes in the range of 2 – 5 (polysome sample) were pooled and treated equally for the remaining purification. The pooled samples were incubated with 100 µl pre-equilibrated magnetic IgG-coupled Dynabeads M-270 Epoxy (Life Technologies) for 1 h at 4 °C. Beads were washed with 3 x 1 ml wash buffer (20 mM HEPES pH 7.4, 100 mM KOAc, 10 mM Mg(OAc)_2_, 1 mM DTT, 10 µg/ml tigecycline). The Ccr4-Not:ribosome complexes were eluted from IgG beads by incubation with 70 U AcTEV protease (Thermo Fisher) in 50 µl elution buffer (20 mM HEPES pH 7.4, 100 mM KOAc, 10 mM Mg(OAc)_2_, 1 mM DTT) for 1.5 h at 4 °C. In the case of the cross-linked sample, chemical cross-linking was performed using 0.5 mM BS3 (Thermo Fisher) for 30 min on ice, before the reaction was quenched with 20 mM Tris. All samples were kept on ice until cryo-EM grid preparation, or were frozen in liquid nitrogen and stored at −80 °C. For northern blotting in presence of cycloheximide, the purification was performed following the same protocol, but instead of tigecycline, 100 µg/ml cycloheximide were used in the lysis buffer and 10 µg/ml cycloheximide during sucrose density centrifugation and affinity purification.

The sample leading to the eIF5A-bound AA/PP/-- ribosome was prepared by culturing the Not4-FTPA strain in 10 l YPD medium supplemented with 5 µg/ml ampicillin and 10 µg/ml tetracycline and harvesting by centrifugation at OD_600_ of 0.9. The pellet was washed in water and lysis buffer (20 mM HEPES pH 7.4, 100 mM KOAc, 10 mM Mg(OAc)_2_, 1 mM DTT, 0.5 mM PMSF, 100 µg/ml cycloheximide, protease inhibitor cocktail tablet (Roche)) and frozen in liquid nitrogen. Lysis was performed by grinding the frozen cell pellet in a Freezer Mill (6970 EFM). The ground powder was thawed in 15 ml lysis buffer and the lysate was cleared by centrifugation at 12000 x g for 15 min at 4 °C. Cleared lysate was incubated with 100 µl pre-equilibrated magnetic IgG-coupled Dynabeads M-270 Epoxy (Life Technologies) for 1 h at 4 °C. Beads were washed with 3 x 2 ml wash buffer (20 mM HEPES pH 7.4, 100 mM KOAc, 10 mM Mg(OAc)_2_, 1 mM DTT, 10 µg/ml cycloheximide). Elution was performed by incubation with 200 U AcTEV protease (Thermo Fisher) in 300 µl elution buffer (20 mM HEPES pH 7.4, 100 mM KOAc, 10 mM Mg(OAc)_2_, 1 mM DTT) for 1.5 h at 4 °C. The eluate was layered on top of a sucrose density gradient (10-50% sucrose in 20 mM HEPES pH 7.4, 100 mM KOAc, 10 mM Mg(OAc)_2_, 1 mM DTT, 10 µg/ml cycloheximide in 14 × 95 mm polyallomer tubes, SW40, Beckman Coulter) and centrifuged at 192072 x g for 2.5 h at 4 °C. The gradient was subsequently fractionated from top to bottom using a Gradient Master (BioComp). The 80S fraction was pelleted for 1 h at 436000 x g at 4 °C (TLA-100, Beckman Coulter), resuspended in 20 mM HEPES pH 7.4, 100 mM KOAc, 10 mM Mg(OAc)_2_, 1 mM DTT, 10 µg/ml cycloheximide and kept on ice until cryo-EM grid preparation.

### Electron microscopy and image processing

For all samples, 0.05% β-octylglucoside was added shortly before grid preparation. Cryo-EM grids were prepared by application of 3.5 µl of the sample containing purified Ccr4-Not:ribosome complexes onto R3/3 copper grids with 3 nm continuous carbon support (Quantifoil) and vitrification in liquid ethane using a Vitrobot Mark IV (Thermo Fisher). Data collection was performed on a Titan Krios TEM (Thermo Fisher) equipped with a Falcon II direct electron detector. Data were collected at 300 kV with a total dose of 25 e^‐^/Å^2^ fractionated over 10 frames with a pixel size of 1.084 Å/pixel and a target defocus range of −1.3 to −2.8 μm using software EPU (Thermo Fisher). The raw movie frames were aligned using MotionCor2 (*32*) and contrast transfer function (CTF) estimation was performed using Gctf (*33*).

For the tigecycline monosome dataset, 9945 micrographs were used for automated particle picking with Gautomatch resulting in 557,059 initial particles, of which 359,890 were selected for further processing upon 2D classification in RELION-2.1 (*34*). After an initial round of 3D refinement, 3D classification with a mask on the ribosomal inter-subunit space was performed. The dataset featured two major classes: a ribosome lacking tRNAs but containing eIF5A (~31% of the data) and a ribosome containing P-site tRNA and additional density in the E-site, which was later identified as N-terminal domain of Not5 (~48% of the data). The 176111 particles belonging to the Not5 containing class were refined to a final resolution of 2.8 Å using 3D refinement, ctf-refinement and post-processing in RELION-3.0 (*35*). The final volume was filtered according to local resolution using RELION-3.0. A classification scheme is provided in fig. S4A.

For the polysome dataset, 9611 micrographs were used for automated particle picking with Gautomatch resulting in 828,516 initial particles, of which 435,002 were selected for further processing upon 2D classification in RELION-3.0. After an initial round of 3D refinement, 3D classification was performed with a mask on the ribosomal inter-subunit space, yielding 3 major classes: a ribosome lacking tRNAs but containing eIF5A (~31% of the data), an AP/EE hybrid (~39% of the data) and a ribosome containing P-site tRNA and Not5 (~21% of the data). The Not5 containing class was further classified using a mask on Not5 and finally without a mask, to further enrich particles containing Not5 and to remove bad particles. The final class containing P-site tRNA, Not5 and density for a nascent chain was refined to a resolution of 3.2 Å using 3D refinement, ctf-refinement and post-processing in RELION-3.0. The final volume was filtered according to local resolution using RELION-3.0. A classification scheme is provided in fig. S4B.

For the cycloheximide monosome dataset 12,697 micrographs were used for automated particle picking with Gautomatch resulting in 1,378,230 initial particles, of which 1,194,480 were selected for further processing upon 2D classification in RELION-3.0. After an initial round of 3D refinement and 3D classification, bad particles and empty ribosomes were discarded, resulting in 274,817 particles (100%) featuring A-, and/or P-site tRNA, which were used for extensive 3D classification. This resulted in a class of AA/PP/-- ribosomes containing eIF5A (~19%), a class of --/PP/--ribosomes containing Not5 (~18%), a class of ribosomes containing P-site tRNA and eRF1 in the A-site (~19%), a class of AA/PP/-- ribosomes in a non-rolled state containing Not5 (~11%), a class of AA/PP/-- ribosomes (~3%), a class of --/PP/-- ribosomes (~16%) and a class of empty ribosomes bound to a hibernation factor (~14%). The 50,989 particles belonging to the eIF5A containing AA/PP/-- class were refined to a final resolution of 3.1 Å using 3D refinement, ctf-refinement and post-processing in RELION-3.0. The final volume was filtered according to local resolution using RELION-3.0. A classification scheme is provided in fig. S4C.

For the cross-linked monosome dataset, 10,654 micrographs were used for automated particle picking with Gautomatch resulting in 589,191 initial particles, of which 337,215 were selected for further processing upon 2D classification in RELION-3.0. After an initial round of 3D refinement, 3D classification was performed using a mask comprising the ribosomal P- and E-sites, yielding a major class, containing P-site tRNA and Not5 (~37% of the data). The class was further subclassified without a mask to remove low quality particles. The final volume was refined to 4.0 Å using RELION-3.0. A classification scheme is provided in fig. S4D.

### Model building and refinement

A full model was built for the Not5-bound --/PP/-- ribosome. The model was based on the crystal structure of the yeast 80S ribosome (PDB code 4V88). Ribosomal proteins and RNA were modeled and refined using COOT (*36*) and PHENIX (*37*). A second copy of the ribosomal protein eL41 was identified adjacent to helix 54, modeled and refined. Initiator-tRNA was modeled based on the crystal structure of yeast tRNA_i_^Met^ (PDB code 1YFG) and refined. Not5^2-113^ was *de-novo* modeled and refined.

The model of the eIF5A-bound AA/PP/-- ribosome was derived from the Not5-bound ribosome. The model of eIF5A and the A-site tRNA were derived from a previously solved cryo-EM structure (PDB code 5GAK). The models were refined to fit the AA/PP/-- density map using COOT and PHENIX.

Cryo-EM densities and molecular models were visualized using ChimeraX (*38*).

### Probe generation for tRNA Northern blotting

Specific digoxigenin (DIG) labeled antisense probes for tRNA Northern blotting were generated by amplifying an unrelated DNA sequence of 230 nt from a pcDNA5/FRT/TO derived plasmid (*39*) by PCR in presence of 1 mM DIG-dUTP. The reverse primer used in the PCR reaction had a 3’-overhang of 30 nt which were reverse complement to the initial 30 bases at the 5’ of the respective tRNAs.

Hybridizing sequence for tRNA_i_^Met^: 5’- CCCTGCGCGCTTCCACTGCGCCACGGCGCT-3’

Hybridizing sequence for tRNA^Ala^: 5’- GGAGCGCGCTACCGACTACGCCACACGCCC-3’

### tRNA Northern blotting

The samples for tRNA northern blotting were prepared as described above. In essence, cells were lyzed in presence of either no antibiotic, 100ug/ml tigecycline, or 100ug/ml cycloheximide. For sucrose density gradient centrifugation, the concentrations of antibiotics were reduced to 10ug/ml. After fractionation, samples of the total 80S fractions of all preparations were taken and the remaining 80S fractions were subjected to affinity purification as described above. Upon elution of the complexes through tag cleavage using AcTEV protease (Thermo Fisher), the ribosome concentrations were measured according to A254.

The northern blots were performed according to Roche DIG Northern Starter Kit protocol. Equal amounts of ribosomes (~160 pmol) of each sample and 100 ng of yeast tRNA mix as control were separated on NOVEX TBE-Urea Gels 10% (Invitrogen), transferred to Nylon Hybond N+ membranes (Amersham) and probed against tRNAs with DIG-labeled antisense probes. Chemo luminescence was detected on an Amersham 600 imager (GE Healthcare).

### Selective ribosome profiling and data analysis

The yeast strain with genomically tagged Not4 (Not4-FTPA) and uL30 (uL30-TAP) were cultured in YPD medium and harvested them by centrifugation at OD_600_ of 0.6. The harvested cell pellet was frozen in liquid nitrogen, and then ground in liquid nitrogen using a mortal. The cell powder was resuspended with no drug lysis buffer (50 mM Tris pH7.5, 100 mM NaCl, 10 mM MgCl_2_, 0.01% NP-40, 1 mM DTT) to prepare the lysate. The lysate was centrifuged at 39,000 g for 30 min at 4 ºC, and the supernatant fraction was used for the purification step. Not4-FTPA or uL30-TAP bound ribosome were affinity purified using IgG conjugated Dynabeads (Invitrogen), followed by TEV protease cleavage to release them.

Library preparation was performed according to the method previously described with following modifications (*40*). TEV eluate containing 20 µg of total RNA was treated with 10 units of RNase I (Epicentre) at 24 ºC for 45 min. As linker DNA, 5’- (Phos)NNNNNIIIIITGATCGGAAGAGCACACGTCT GAA(ddC)-3’ where (Phos) indicates 5’ phosphorylation and (ddC) indicates a terminal 2’, 3’-dideoxycytidine, was used. The Ns and Is indicate a random barcode for eliminating PCR duplication and multiplexing barcode, respectively. The linkers were pre-adenylated with a 5’ DNA Adenylation kit (NEB), and then used for the ligation reaction. Un-reacted linkers were digested by 5’ deadenylase (NEB) and RecJ exonuclease (epicentre) at 30°C for 45 min. An oligo 5’- (Phos)NNAGATCGGAAGAGCGTCGTGTAGGGAA AGAG(iSp18)GTGACTGGAGTTCAGACGTGTGCT C-3’, where (Phos) indicated 5’ phosphorylation and Ns indicate a random barcode, was used for reverse transcription. PCR was performed with oligos 5’- AATGATACGGCGACCACCGAGATCTACACTCTTTCCCTACACGACGCTC-3’ and 5’- CAAGCAGAAGACGGCATACGAGATJJJJJJGTGAC TGGAGTTCAGACGTGTG-3’, where Js indicate the reverse complement of the index sequence discovered during Illumina sequencing. The libraries were sequenced on a HiSeq 4000 (Illumina).

Reads were mapped to yeast transcriptome, removing duplicated reads based on random barcode sequences. The analysis of Not4- and uL30-selective ribosome profiling were restricted to 21, 22, 28, 29, 30 nt and 21, 22, 23, 29, 30, 31 nt long reads, respectively. We estimated the position of the A site from the 5’-end of the reads at the initiation codon based on the length of each footprint. The offsets were 17 for 23 nt reads and 16 for 22, 29, 30, and 31 nt reads, and 15 for 21 and 28 nt reads.

Ribosome pause scores were calculated as previously published (*41*). Basically, we took the footprints accumulation over the average of the footprint density in the given ORFs. Analyses were restricted to the mRNAs with 0.5 footprints/codon and more. The averaged pause scores on given codons were computed.

### Analytical sucrose density gradient centrifugation

Yeast cells were grown exponentially at 30 ºC and treated 0.1 mg/ml of cycloheximide for 5 min before harvesting, then harvested by centrifugation. The harvested cell pellet was frozen and ground in liquid nitrogen using a mortal. The cell powder was resuspended with lysis buffer (20 mM HEPES-KOH, pH 7.4, 100 mM potassium acetate, 2 mM magnesium acetate, 0.5mM dithiothreitol, 1mM phenylmethylsulfonyl fluoride, 1tablet/10ml Complete mini EDTA-free (#11836170001, Roche)) to prepare the lysate. The lysate (the equivalent of 50 A260 units) were layered on top of the 10 – 50 % sucrose gradients and then centrifuged at 150,000×g in a P28S rotor (Hitachi Koki, Japan) for 2.5 hr at 4ºC. The polysome profiles were generated by continuous absorbance measurement at 254 nm using a single path UV-1 optical unit (ATTO Biomini UV-monitor) connected to a chart recorder (ATTO digital mini-recorder). Proteins in each fraction was separated by 10% Nu-PAGE, and were transferred to PVDF membranes (Millipore; IPVH00010). After blocking with 5 % skim milk, the blots were incubated with the Anti-HA-Peroxidase (#12013819001, Roche), and detected by ImageQuant LAS4000 (GE Healthcare).

### Transcriptional shut-off and RNA Northern blot analysis

For the *GAL1* UAS transcriptional shut-off analysis, cells expressing the appropriate plasmids were grown at 24 °C in synthetic media with 2% galactose/1% sucrose to allow for expression of the reporter mRNA. Cells were shifted to synthetic media without sugar at an OD600=0.4, and then transcription was repressed by adding glucose to a final concentration of 4%. Cells were collected at the time points indicated in the figures. For the cycloheximide (CHX) treated experiments, different concentrations of CHX (0, 0.2, 50 or 100 mg/ml) were added to cells (OD600=0.4) for 40 min followed by the transcriptional shutoff analysis.

Total RNA was extracted by phenol/chloroform and precipitated with 95% EtOH overnight. 30μg of RNA was separated on 1.4% agarose-formaldehyde gels at 100 V for 1.5 hr, transferred to nylon membranes, and probed with 32P-labeled antisense oligonucleotides to detect poly (G) (oJC168), *HIS3* (oJC2564), or *SCR1* (oJC306) (Table S2). Blots were exposed to PhosphorImager screens, scanned by Typhon 9400, and quantified by ImageQuant.

### Polyribosome analysis and Western blotting

100 μg/mL CHX were added when cells reach to OD600=0.4 before harvesting. For detecting Dhh1p by western blotting, cells were crosslinked at a final concentration of 0.25% formaldehyde for 5 min, then treated with 125 mM glycine for 10 min to quench crosslinking before adding CHX. Cells were then lysed into lysis buffer (10 mM Tris pH 7.4, 100 mM NaCl, 30 mM MgCl2, 1 mM DTT, 100 μg/Ml CHX) by vortexing with glass beads, and cleared using the hot needle puncture method. 1% Triton X-100 was added into the supernatant. 7.5 OD260 units were loaded on 15%–45% (w/w) sucrose gradients prepared on a Biocomp Gradient Master in gradient buffer (50 mM Tris-acetate pH 7.0, 50 mM NH4Cl, 12 mM MgCl2, 1 mM DTT) and centrifuged in a SW-41Ti rotor at 41,000 rpm for 2 hr and 26 min (In order to detect Dhh1p signal, the centrifugation step is optimized to 1 hr and 13 min for all groups in Fig. 5F (*42*) at 4 °C. Gradients were fractionated by using a Brandel Fractionation System and an ISCO UA-6 ultraviolet detector. Fractions were precipitated at −20 °C with 10% TCA (final concentration) overnight. Pellets were washed with 80% acetone, resuspended in 50 ul of SDS-PAGE loading buffer, and boiled at 95 °C for 5 min, then separated by 10% SDS polyacrylamide gels, followed by Western blotting with primary antibodies (anti-HA [BioLegend, PRB-101C], anti-Rpl4 [Proteintech, 11302-1-AP], anti-Dhh1p at 4 °C overnight and incubated with secondary antibodies (goat-anti-Mouse [Santa Cruz sc-2005] and goat-anti-Rabbit [Pierce 31460]) at room temperature for 1 hr. Signal was detected by chemiluminescence using Blue Ultra Autorad film.

## Acknowledgments

We thank Dr. Shintaro Iwasaki, Manabu Ishii, Itoshi Nikaido and the bioinformatics analysis environment service on RIKEN cloud at RIKEN advanced center for computing and communication for support in computational analyses and Heidemarie Sieber, Charlotte Ungewickell, Susanne Rieder, Lukas Kater and Jonathan Schneider for technical support. This work used the Vincent J. Coates Genomics Sequencing Laboratory at UC Berkeley, supported by NIH S10 Instrumentation Grants OD018174 Instrumentation Grant. We also thank Dr. Thomas Fröhlich and LAFUGA for mass-spectrometry analysis.

## Author contributions

Experiments were designed by R.Bu., R.Be., T.I. and J.Co.; O.B. collected cryo-EM data; R.Bu. prepared cryo-EM samples and processed the cryo-EM data; R.Bu. and J.Ch. built molecular models and R.Bu. and T.B. analyzed the structures; Yo.M. performed and analyzed selective ribosome profiling; YH.C. and Ya.M. performed analytical ultracentrifugation; A.G. and R.Bu. performed tRNA Northern blotting; YH.C., N.A., Th.S., Ta.S. and K.I. performed mRNA stability measurements; the manuscript was written by R.Bu., R.Be. and J.Co.; all authors discussed the results and commented on the manuscript.

## Data availability

The sequencing data for ribosome profiling experiments will be deposited in NCBI’s Gene Expression Omnibus and will be accessible through GEO series accession number …. Cryo-EM maps and respective molecular models will be deposited with accession IDs EMD-… and PDB-….

## Supplementary Materials

Figures S1-S4

Tables S1-S2

References (*43, 44*)

