## Supplementary Data for "The Ccr4-Not complex monitors the translating ribosome for codon optimality"

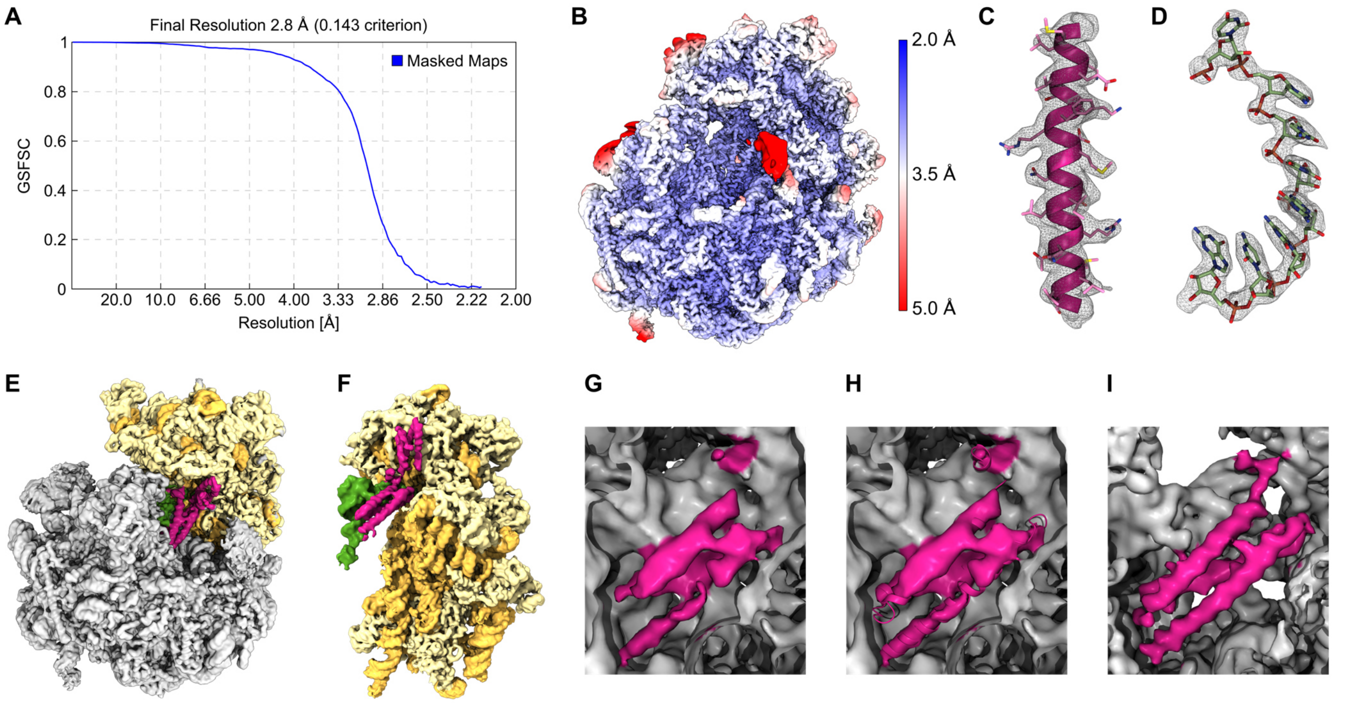


**Fig. S1.** (**A**) Gold Standard Fourier shell correlation of the Not5 bound 80S ribosome (**B**) Cryo-EM density of the Not5 bound 80S ribosome colored according to local resolution. (**C**) Representative density of the Not5-NTD. (**D**) Representative density of ribosomal RNA. (**E**) Cryo-EM density of the cross-linked Not5 bound 80S ribosome (**F**) Cryo-EM density of the 40S subunit, the P-site tRNA and the extended Not5-NTD of the cross-linked Not5 bound bound 80S ribosome. (**G**) Cryo-EM density of the Not5 bound 80S ribosome in absence of antibiotics. (**H**) Cryo-EM density of the Not5 bound 80S ribosome in absence of antibiotics with a rigid body docked model of Not5-NTD. (**I**) Cryo-EM density of the Not5 bound 80S ribosome stabilized by tigecycline.

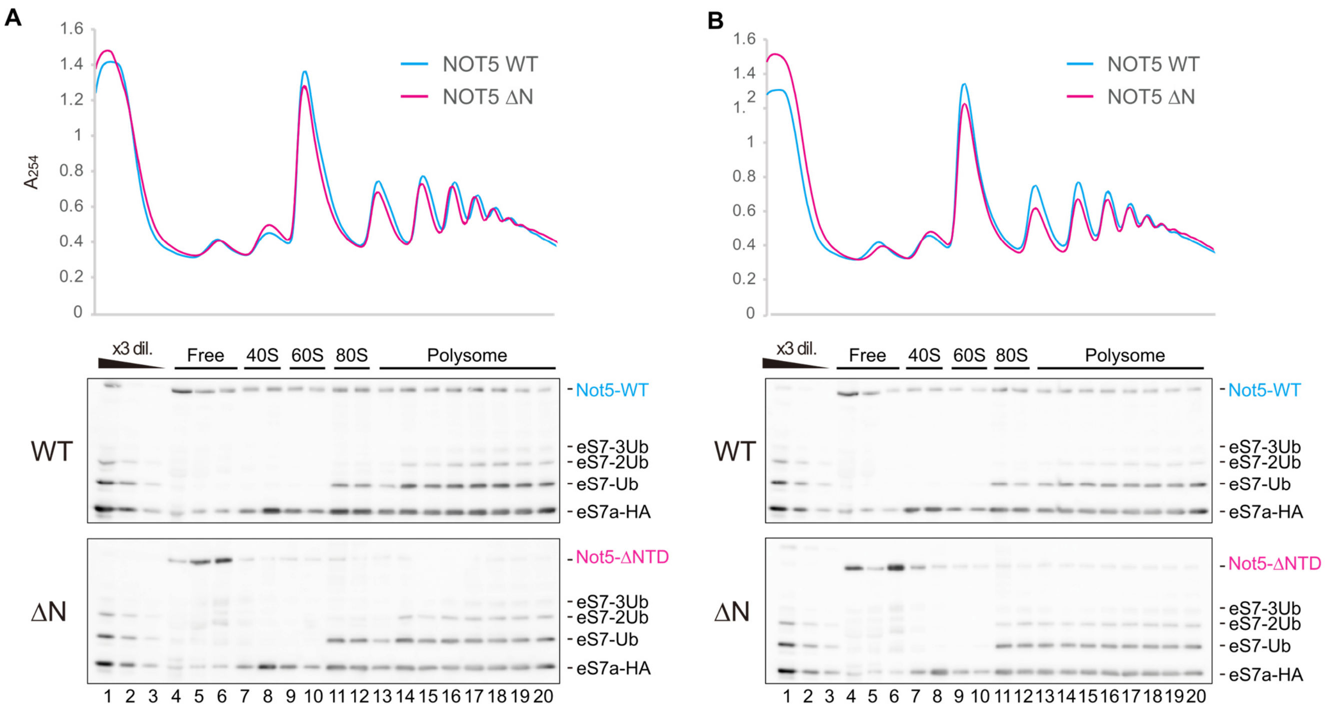


**Fig. S2.** (**A** to **B**). Replicates of density gradient fractionation profiles of Not5-HA and Not5-ΔNTD-HA strains and corresponding western blot detection of the respective Not5 proteins and of ribosomal protein eS7-HA. The signals at higher molecular weight correspond to ubiquitinated species of eS7-HA.

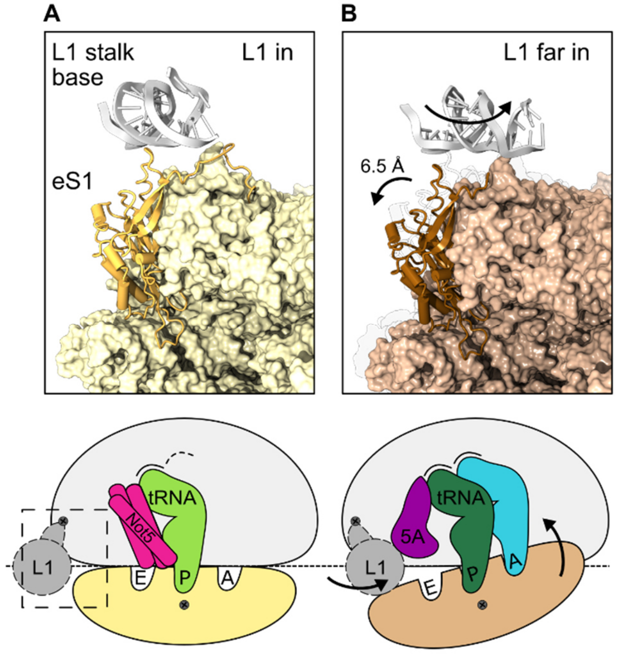


**Fig. S3.** (**A**) In-position of the L1 stalk base in the non-rolled --/PP/-- ribosome. (**B**) Far in-position of the L1 stalk base in the eIF5A bound rolled AA/PP/-- ribosome.

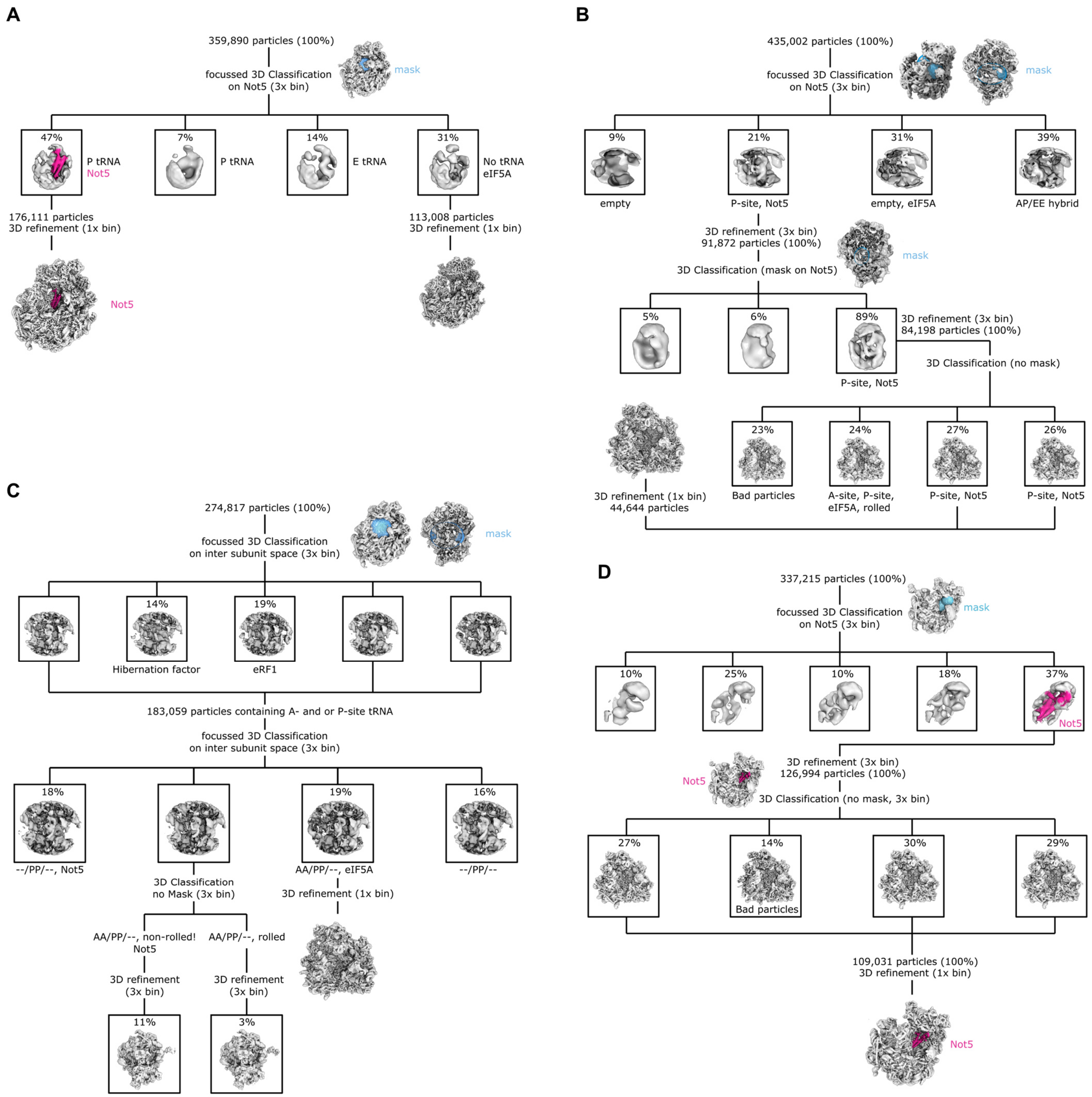


**Fig. S4.** (**A**) Sorting scheme of the tigecycline stalled monosome sample. (**B**) Sorting scheme of the tigecycline stalled polysome sample. (**C**) Sorting scheme of the cycloheximide stalled monosome sample. (**D**) Sorting scheme of the tigecycline stalled cross-linked monosome sample.

**Table S1. Yeast strains used in this study**

| Name | Genotype | Source |
|---|---|---|
| yJC151 | MATa, ura3, leu2, his3, met15 | (*43*) |
| yJC1708 | ura3, leu2, his3, met15 or MET15, LYS2, not5::LEU2 | This study |
| yJC1892 | MATa, ura3, leu2, his3, met15 [pGAL-PGK1-SYNOP-pG, URA3] | (*5*) |
| yJC1893 | MATa, ura3, leu2, his3, met15 [pGAL-PGK1-SYNNONOP-pG, URA3] | (*5*) |
| yJC1913 | MATa, ura3, leu2, his3, met15, dhh1::NEO [pGAL-PGK1-SYNOP-pG, URA3] | (*5*) |
| yJC1914 | MATa, ura3, leu2, his3, met15, dhh1::NEO [pGAL-PGK1-SYNNONOP-pG, URA3] | (*5*) |
| yJC1917 | MATa, ura3, leu2, his3, lys2, dcp2::NEO [pGAL-PGK1-SYNOP-pG, URA3] | (*5*) |
| yJC1918 | MATa, ura3, leu2, his3, lys2, dcp2::NEO [pGAL-PGK1-SYNNONOP-pG, URA3] | (*5*) |
| yJC1961 | MATa, ura3, his3, leu2, met15, ccr4:NEO [pGAL-PGK1-SYNOP-pG, URA3] | (*5*) |
| yJC1962 | MATa, ura3, leu2, his3, met15, ccr4::NEO [pGAL-PGK1-SYNNONOP-pG, URA3] | (*5*) |
| yJC2364 | MATa, ura3, his3, leu2, met15, pop2::NEO [pGAL-PGK1-SYNOP-pG, URA3] | (*5*) |
| yJC2365 | MATa, ura3, his3, leu2, met15, pop2::NEO [pGAL-PGK1-SYNNONOP-pG, URA3] | (*5*) |
| yJC2499 | MATa, ura3, leu2, his3, met15, [N-terminally optimal FLAG tagged 0% optimal HIS3 under the control of the GAL1 promoter, URA3] | (*7*) |
| yJC2504 | MATa, ura3, leu2, his3, met15, [N-terminally optimal FLAG tagged 50% optimal HIS3 under the control of the GAL1 promoter, URA3] | (*7*) |
| yJC2509 | MATa, ura3, leu2, his3, met15, [N-terminally optimal FLAG tagged 100% optimal HIS3 under the control of the GAL1 promoter, URA3] | (*7*) |
| yJC2587 | ura3, leu2, his3, met15 or MET15, LYS2, not5::LEU2, [pGAL-PGK1-SYNOP-pG, URA3] | This study |
| yJC2588 | ura3, leu2, his3, met15 or MET15, LYS2, not5::LEU2, [pGAL-PGK1-SYNNONOP-pG, URA3] | This study |

| | | |
|---|---|---|
| yJC2708 | MATa, ura3, leu2, his3, met15, NOT5-HA::HIS | This study |
| yJC2721 | ura3, leu2, his3, met15 or MET15, LYS2, not5::LEU2, [N-terminally optimal FLAG tagged 0% optimal HIS3 under the control of the GAL1 promoter, URA3] | This study |
| yJC2722 | ura3, leu2, his3, met15 or MET15, LYS2, not5::LEU2, [N-terminally optimal FLAG tagged 50% optimal HIS3 under the control of the GAL1 promoter, URA3] | This study |
| yJC2723 | ura3, leu2, his3, met15 or MET15, LYS2, not5::LEU2, [N-terminally optimal FLAG tagged 100% optimal HIS3 under the control of the GAL1 promoter, URA3] | This study |
| yJC2809 | MATa, ura3, leu2, his3, lys2, rps25a::KANMX, rps25b::KANMX [pGAL-PGK1-SYNOP-pG, URA3] | This study |
| yJC2810 | MATa, ura3, leu2, his3, lys2, rps25a::KANMX, rps25b::KANMX [pGAL-PGK1-SYNOP-pG, URA3] | This study |
| W303-1a | MATa ade2 his3 leu2 trp1 ura3 can1 | Lab. Stock |
| Not4-FTPA | MATa ade2 his3 leu2 trp1 ura3 can1 NOT4-FTPA::natNT2 | (*44*) |
| uL30-TAP | MATa ade2 his3 leu2 trp1 ura3 can1 uL30-TAP::TRP1 | (*44*) |

**Table S2. Plasmids and oligos used in this study**

| Name | Description | Source |
|---|---|---|
| pJC296 | PGK1pG reporter (under control of GAL1 UAS) | (*43*) |
| pJC672 | PGK1pG reporter with SYNOP ORF (under control of GAL1 UAS) | (*5*) |
| pJC673 | PGK1pG reporter with SYNNONOP ORF (under control of GAL1 UAS) | (*5*) |
| pJC857 | 0% optimal HIS3 with N-terminal FLAG tag (GAL1 promoter) | (*7*) |
| pJC862 | 50% optimal HIS3 with N-terminal FLAG tag (GAL1 promoter) | (*7*) |
| pJC867 | 100% optimal HIS3 with N-terminal FLAG tag (GAL1 promoter) | (*7*) |
| pRS415-Not5-HA | CEN, LEU2, Not5-HA (NOT5 promoter) | This study |
| pRS415-Not5ΔNTD-HA | CEN, LEU2, Not5ΔNTD-HA (NOT5 promoter), the region from 2 to 113 A.A. is deleted | This study |
| pRS416-eS7-HA | CEN, URA3, eS7a-HA (eS7A promoter) | (*11*) |
| Oligo oJC168 | 5'-AATTCCCCCCCCCCCCCCCCCA-3' | (*43*) |
| Oligo oJC306 | 5'-GTCTAGCCGCGAGGAAGG-3' | (*43*) |
| Oligo oJC2564 | 5'-CCTGATCCAAACCTTTTTACTCC-3' | (7) |
